# Sinusoidal voltage protocols for rapid characterisation of ion channel kinetics

**DOI:** 10.1101/100677

**Authors:** Kylie A. Beattie, Adam P. Hill, Rémi Bardenet, Yi Cui, Jamie I. Vandenberg, David J. Gavaghan, Teun P. de Boer, Gary R. Mirams

## Abstract

Understanding the roles of ion currents is crucial to predict the action of pharmaceuticals and mutations in different scenarios, and thereby to guide clinical interventions in the heart, brain and other electrophysiological systems. Our ability to predict how ion currents contribute to cellular electrophysiology is in turn critically dependent on our characterisation of ion channel kinetics — the voltage-dependent rates of transition between open, closed and inactivated channel states. We present a new method for rapidly exploring and characterising ion channel kinetics, applying it to the hERG potassium channel as an example, with the aim of generating a quantitatively predictive representation of the ion current. We fit a mathematical model to currents evoked by a novel 8 second sinusoidal voltage clamp in CHO cells over-expressing hERG1a. The model is then used to predict over 5 minutes of recordings in the same cell in response to further protocols: a series of traditional square step voltage clamps, and also a novel voltage clamp comprised of a collection of physiologically-relevant action potentials. We demonstrate that we can make predictive cell-specific models that outperform the use of averaged data from a number of different cells, and thereby examine which changes in gating are responsible for cell-cell variability in current kinetics. Our technique allows rapid collection of consistent and high quality data, from single cells, and produces more predictive mathematical ion channel models than traditional approaches.

**Table of Contents Category:** Techniques for Physiology^1^

**Key Points:**

- Ion current kinetics are commonly represented by current-voltage relationships, time-constant voltage relationships, and subsequently mathematical models fitted to these. These experiments take substantial time which means they are rarely performed in the same cell.
- Rather than traditional square-wave voltage clamps, we fit a model to the current evoked by a novel sum-of-sinusoids voltage clamp that is only 8 seconds long.
- Short protocols that can be performed multiple times within a single cell will offer many new opportunities to measure how ion current kinetics are affected by changing conditions.
- The new model predicts the current under traditional square-wave protocols well, with better predictions of underlying currents than literature models. The current under a novel physiologically-relevant series of action potential clamps is predicted extremely well.
- The short sinusoidal protocols allow a model to be fully fitted to individual cells, allowing us to examine cell-cell variability in current kinetics for the first time.

## 1 Introduction

Mathematical models of ion channels are a quantitative expression of our understanding of ion channel kinetics: they express the probability of channels existing in different conformational states (typically, closed, open and inactivated) and the rates of transition between these states (Bett et al., 2011; Vandenberg et al., 2012). Parameterising/calibrating a mathematical model of an ion current is a concise way to characterise ion channel kinetics, to capture our understanding in a quantitative framework, and to communicate this knowledge to others. There have been some notable advances in deriving mathematical models for ion channel behaviour (Balser et al., 1990; Cannon and D’Alessandro, 2006; Siekmann et al., 2011, 2012; Loewe et al., 2015), with some stressing the need for validation/testing of the model using data from the same cell (Tomaiuolo et al., 2012). In this paper we present a new approach for characterising ion channel kinetics, using novel short protocols and parameter inference techniques to construct an ion channel model.

The *KCNH2* gene (also known as *hERG*) has been shown to encode the primary subunit of the voltage-gated ion channel Kv11.1 that carries the rapid delayed rectifier potassium current (I_Kr_) (Trudeau et al., 1995; Sanguinetti et al., 1995). In this article we focus on mathematical modelling of hERG channel kinetics, demonstrating our approach by constructing an improved model of this ion current. hERG plays important roles in the brain (Babcock and Li, 2013); gastrointestinal tract (Farrelly et al., 2003); uterine contractions (Parkington et al., 2014); cell-proliferation and apoptosis (Jehle et al., 2011) and cancer progression (Lastraioli et al., 2015), but I_Kr_ is best known as a repolarising cardiac ion current. The channel is susceptible to binding and blockade by pharmaceutical compounds, which is strongly linked to many cases of drug-induced pro-arrhythmic risk (Redfern et al., 2003; Pollard et al., 2010). Mathematical modelling of cardiac electrophysiology, including I_Kr_, forms a core part of a new proposal for routine *in vitro* and *in silico* safety assessment to replace a human clinical drug safety study (Sager et al., 2014; Fermini et al., 2016). A wide range of different mathematical models have been proposed to describe I_Kr_ (literature models are listed in Appendix A, Table A1).

Fig 1 shows predicted I_Kr_ under three different voltage clamps for 29 literature models. These models were developed to describe different species, cell types, temperatures and isoforms, so variation is expected. In Figure 1B–E each row highlights models developed to represent the same species, cell type and temperature; even models for the same conditions provide highly variable predictions.

**Figure 1:**
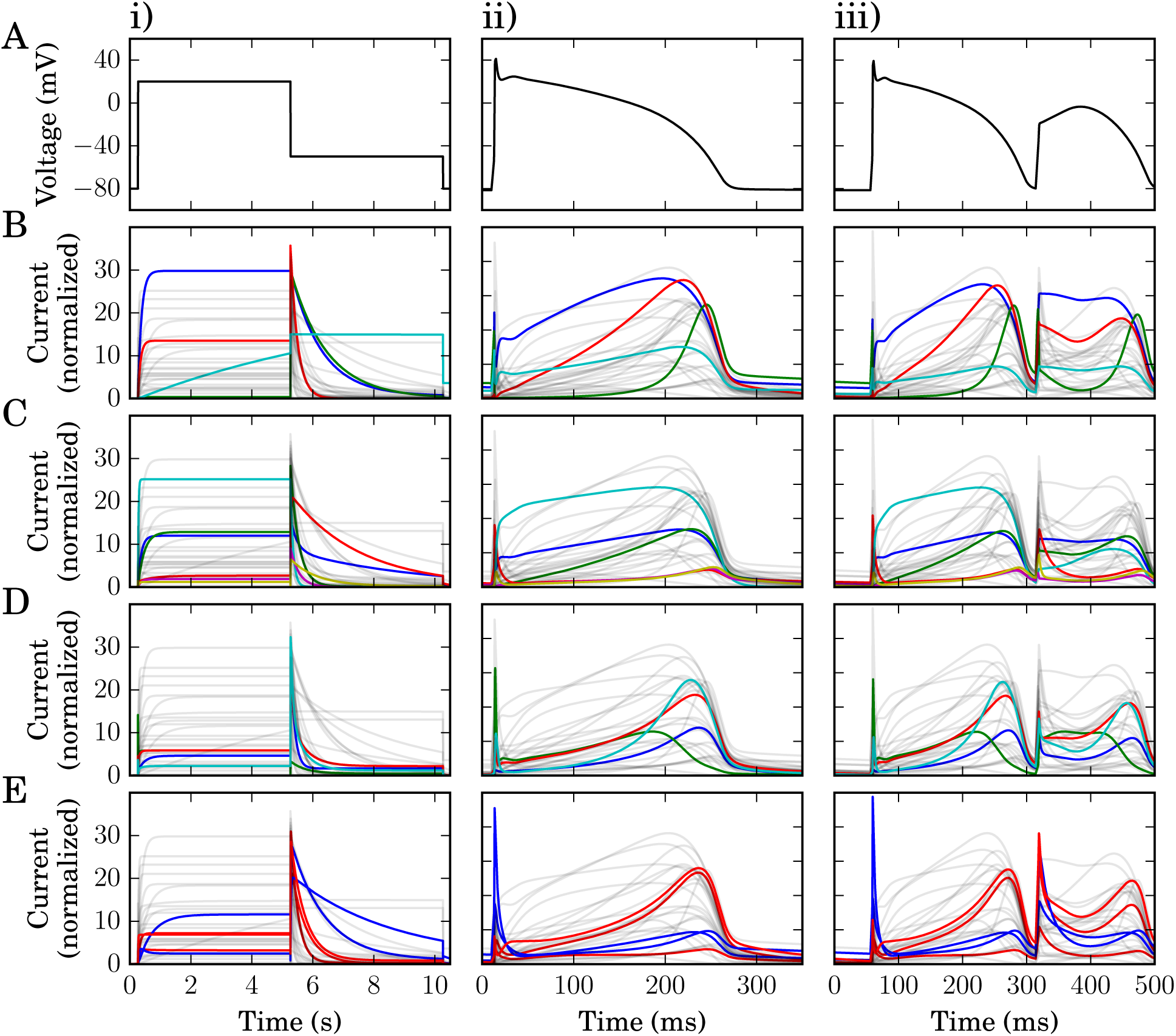
Current predictions from literature models of I_Kr_. Each column shows simulated current predictions from 29 I_Kr_ literature models in response to the different voltage clamp protocols shown in the top row. (**A**) voltage clamps: i) a voltage-step; ii) an action potential; and iii) an action potential displaying pathological properties. Each of the panels below features all 29 current predictions in faint grey, to aid comparison between plots. (**B**) In row B we highlight the four models for canine ventricle at physiological temperature. **C**) In row C we highlight the six models for human ventricle at physiological temperature. (**D**) In row D we highlight the four models for rabbit sino-atrial node at physiological temperature. (**E**) In row E we highlight the five models for hERG1a expression systems: at room temperature in blue; and physiological temperature in red. Currents are normalised such that the maximal conductance is equal to one; i.e. we plot the open probability multiplied by the driving voltage (all model references and structures are listed in Table A1 in Appendix A). All models have been simulated with their original published parameters, with the same reversal potential of –88.4 mV.

The first models of ion channel kinetics were proposed by Hodgkin and Huxley (1952), and relatively little has changed in the methods used for construction of mathematical models of ion channel gating since the original seminal work in this journal in 1952. Their (now traditional) approach is to fit peak currents and time constants of current activation/decay after clamping to fixed voltages; to assemble current/voltage (IV) and time-constant/voltage (*τ*–V) curves; and to describe these curves with interpolating functions.

Condensed voltage clamp step protocols have been suggested as the basis of optimised experiments that provide information about ion channel kinetics faster than experiments to construct IV curves (Hobbs and Hooper, 2008; Fink and Noble, 2009); and optimised current and square step voltage clamps have been used to optimise the fitting of maximal conductances in action potential models (Groenendaal et al., 2015). Single sinusoid voltage clamps have been previously been explored for choosing between possible Shaker channel models that were parameterised using traditional square step voltage clamps (Kargol et al., 2004). Wavelet-based voltage protocols have also been suggested for examining sodium channel dynamics (Hosein-Sooklal and Kargol, 2002). The study by Kargol (2013) features excellent insight into the problem of models behaving similarly under traditional clamps but differently under optimised information-rich protocols. In that paper, these wavelet-based protocols were designed and used to select between Shaker potassium channel models.

In this study, we extend these ideas and propose an 8 second sum-of-sinusoids-based voltage clamp, designed to both explore and fully-characterise the kinetics of the hERG potassium channel. We use this new protocol to record currents from Chinese Hamster Ovary (CHO) cells that are over-expressing hERG1a. These recordings are then used to parameterise a mathematical model which becomes our characterisation of the ion current. We then evaluate the model by predicting the response to both standard square step voltage-clamp protocols and perhaps more importantly physiologically-relevant action potential voltage clamps: using these data (which are independent of the recordings used to fit the model) to perform an extremely thorough validation for the model of ion channel kinetics. Our approach uses a substantially shorter experimental recording to construct the model than the usual approach, which is based on time constants and peak currents from a long series of square step voltage-clamp protocols. As a consequence of the high information content of the short protocol, we are able to generate cell-specific models that advance our understanding of variability of ion currents between cells. Our methodology will be applicable to many ion channels, both in the heart and other electrophysiological systems.

## 2 Methods

### 2.1 Experimental methods

We performed whole-cell patch-clamp voltage clamp experiments, using CHO cells stably expressing hERG1a (Kv11.1) at room temperature. Full details including cell culture, solutions, and equipment settings can be found in Appendix B. In Figure 2 we provide an overview of the experimental approach, denoting the sequence of voltage clamp protocols we performed as Pr0–Pr7.

**Figure 2:**
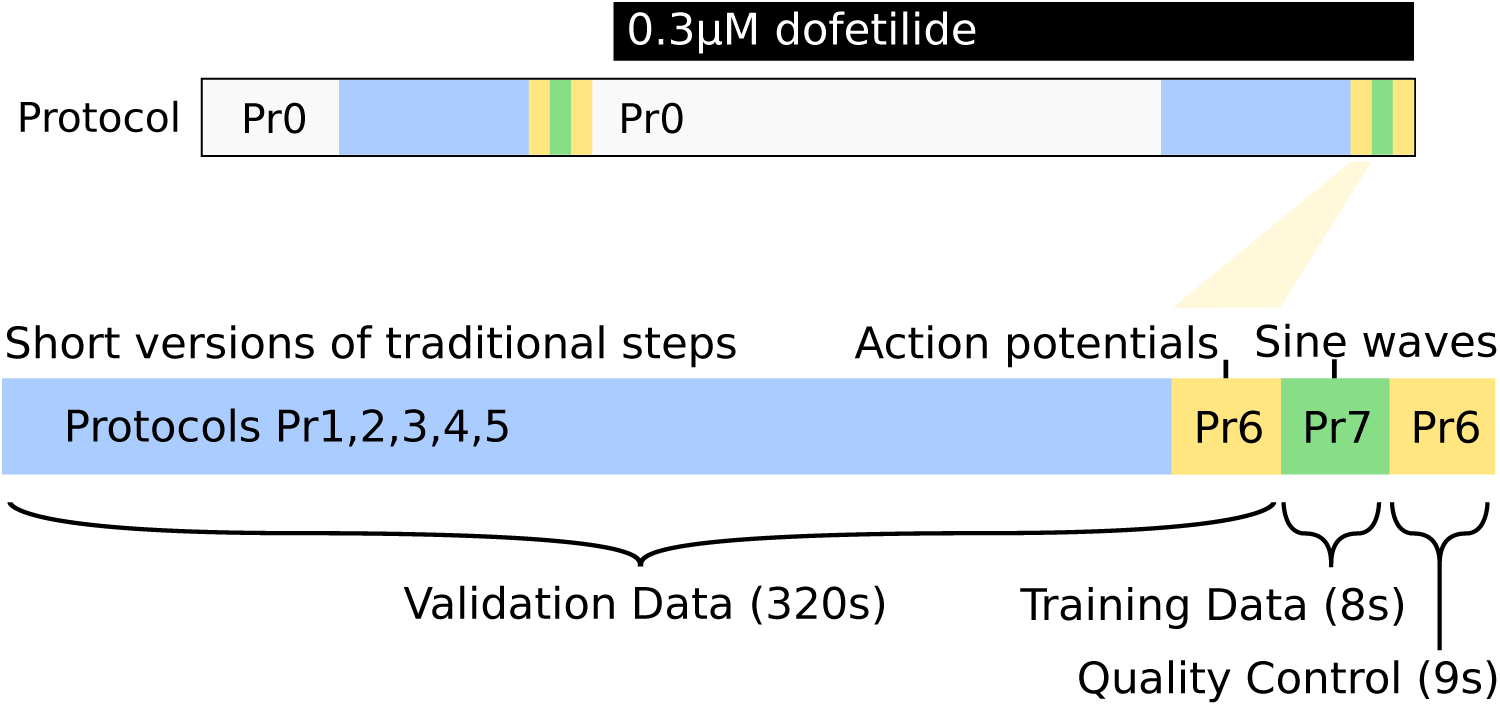
Schematic diagram of the experimental procedure used in this study (not to scale). A simple activation step protocol is repeated in the sections marked ‘Pr0’, before moving on to the highlighted section (below) where data used in the study were recorded. The recording protocols ‘Pr1–7’ are performed twice, once before dofetilide addition, and once after, with the hERG current isolated by subtraction. For full details of the protocols please refer to Appendix B1.4.

In each cell we recorded: a series of conventional voltage-step protocols designed to explore activation (Pr1–3), inactivation (Pr4) and deactivation (Pr5); a new protocol composed of a series of action potential clamps (Pr6 — formed of simulated action potentials from different mathematical models to represent diverse species and pacing frequencies in both healthy and repolarisation-failure conditions); and our new 8s sinusoidal voltage protocol (Pr7, shown in Figure 3). These protocols are all performed in a single experiment using a single cell, and the process can be repeated in different cells. A mathematical model is then fitted/calibrated to solely the current provoked by the sinusoidal protocol, and this model then represents a full characterisation of I_Kr_ in each particular cell. The characterisation is then tested for accuracy by using the fitted mathematical model to predict the results of all the other voltage clamp protocols performed in that cell. Full details of all protocols are given in Appendix B1.4.

**Figure 3:**
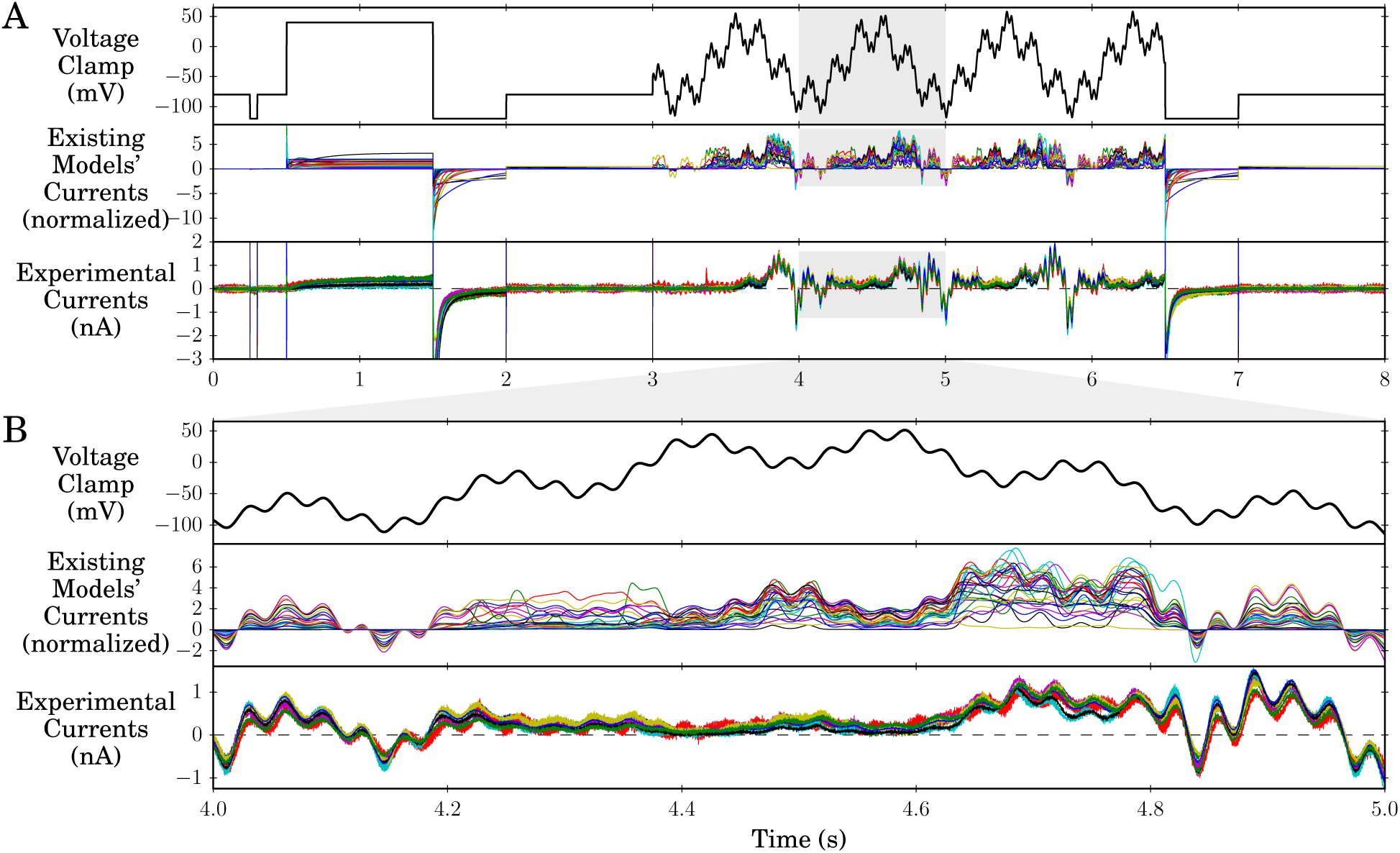
The sinusoidal protocol and example recordings. **A: Top row:** The full sinusoidal voltage protocol (Pr7). **Middle row**: Simulations of expected behaviour in response to this protocol from existing I_Kr_ and hERG models, normalised by scaling the conductance value for each model to minimise the absolute difference between each trace and a reference trace. For calculation of the reversal potential, a temperature of 21.5 °C was used to match the mean experimental conditions. **Bottom row**: Raw data (following leak and dofetilide subtraction) from experimental repeats at room temperature from 9 cells. Experimental traces have been scaled, to remove the effect of different maximal conductances, by a factor chosen to minimise the absolute differences between each trace and a reference experimental trace (that with the peak current during the sinusoidal portion of Pr7). **B**: an enlargement of the highlighted sections of panel A. Whilst there is some variation between cells in the experimental results, they are much more consistent than the predictions from the different models.

In all protocols, the holding potential was initially –80 mV before applying a 50 ms leak step to –120mV before returning back to –80 mV, with this step being used to estimate leak current (as described below in Section 2.2). A voltage step to –120mV at the end of all the protocols ensures that channels close quickly, reducing the time needed between protocols to regain a steady closed state.

#### Protocols 0 to 5 — Square Step Clamps

Protocol 0 is a simple repeated activation pulse designed to open the channel to visually test the recordings were stable and to allow dofetilide binding, considered open state dependent, to occur (see Section 2.3, below). This current was not recorded or used in the subsequent analysis (hence ‘Protocol 0’).

Protocols 1–5 are adaptations of ‘traditional’ square step voltage clamps used in previous studies to examine activation (Pr1–3), inactivation (Pr4) and deactivation (Pr5). Details of the protocol voltages and timings can be found in Appendix B1.4.

The ‘adaptation’ is that protocols 1–5 are shorter than those previously used to calibrate mathematical models (as in fewer test voltages/timings are used), so that it is possible to perform them all in a single cell, with and without dofetilide subtraction. A ‘traditional approach’ would take longer than the experiments performed here, generally requiring multiple cells.

#### Protocol 6 — Action Potentials Clamp

This protocol was formed by combining a series of different simulated action potentials from the Cardiac Electrophysiology Web Lab (Cooper et al., 2016). The range of models we used for the simulations encompassed different cell types, species, and pacing rates. We also added some simulated action potentials where early or delayed after-depolarisations had been induced, to test I_Kr_ behaviour in pro-arrhythmic or pathological settings. The action potentials were shifted slightly so that their resting potentials were exactly –80 mV (see the Supplementary Code for full details and code to reproduce this protocol).

#### Protocol 7 — Sinusoidal Clamp

The protocol used to characterise the current and train the model is a voltage clamp comprised of simple steps and a main sinusoidal section that is in the form of a sum of three sine waves of different amplitudes and frequencies, designed to rapidly explore hERG channel kinetics. The underlying rationale is to force the protocol to ‘sweep’ both the time and voltage dependence of the current gating over physiological voltage ranges.

The start of the protocol takes the form of a leak step followed by a simple activation step which is similar to Protocol 0. This activation step was included to improve the identifiability of the maximal conductance parameter (as described in Appendix B2.2) after preliminary experiments suggested this might improve what is known as ‘parameter identifiability’ (to pin down possible values of the parameter more accurately, and prevent other kinetic parameters compensating for an inaccurate conductance value).

The main sinusoidal portion of the protocol takes the form of a sum of three sine waves as shown in Equation (1):

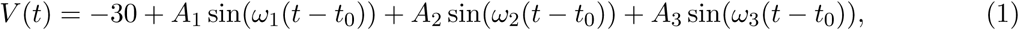

where *A*_1_ = 54 mV, *A*_2_ = 26 mV, *A*_3_ = 10 mV, *ω*_1_ = 0.007 ms^−1^, *ω*_2_ = 0.037 ms^−1^ and *ω*_3_ = 0.19 ms^−1^, and *t* is time measured in milliseconds.

In terms of frequencies, existing models and I_Kr_ recordings include characteristic timescales of order 10 ms to one second (Wang et al., 1997; Zhou et al., 1998). Therefore we designed the sinusoidal protocol’s three frequencies to probe channel kinetics across all these orders of magnitude (10 ms, 100ms and 1 s timescales). We selected frequencies that were co-prime rather than exactly multiples of ten: *ω*_1_ to *ω*_3_ are ordered slow to fast and correspond approximately to sine waves of period 900, 170 and 33 ms, respectively. The aim was that the three distinct frequencies should not become ‘in phase’: the protocol never repeats patterns that the cell has experienced before (ensuring new information is supplied throughout). The offset *t*_0_ is 2500ms as explained in Appendix B1.4. If one was to study other ion channels, these frequencies may need adjustment to examine relevant timescales.

To decide the amplitudes, the oscillations are centred around –30 mV so that a physiological range is explored (–120 < *V* < 60mV). The amplitudes of the sine waves were selected to keep the protocol within this range (*A*_1_ + *A*_2_ + *A*_3_ = 90 mV) and to ensure that *A*_1_ > *A*_2_ > *A*_3_ so that the fastest timescale had the smallest oscillations (to avoid the faster gating processes masking the voltage-dependence of slower ones).

A key step in settling on this particular protocol was its performance in *synthetic data* studies. In these studies we simulated I_Kr_ with different sets of given parameters, then attempted to recover these parameters blindly — using just the generated current trace with added noise, as illustrated in Appendix C2 (we also show this for an I_Ks_ model with the same protocol in Appendix G).

The sinusoidal protocol is of only eight seconds duration, which enables efficient data collection, with training and validation data collected from the same cell. In Figure 3 we present the novel sinusoidal protocol Pr7, the simulated predicted currents from existing models, and the currents we recorded experimentally. The new protocol provokes an even wider array of different behaviours from the existing literature I_Kr_ models (middle panels in Figure 3) than the existing voltage step or action potential clamps (Figure 1); even among models constructed in/for similar conditions/species.

### 2.2 Leak Corrections

We used the leak-step from –80 mV to –120 mV in order to leak-correct the experimental data, according to:

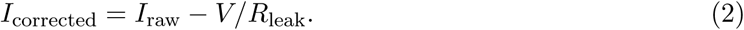

We identified the most appropriate *R*_leak_ value to minimise the difference between the mean current value during the leak step (to –120 mV) compared to the mean value at a holding potential of –80 mV, whilst ensuring that the trace was not over-corrected (which would result in negative currents during the initial stages of activation).

We manually selected leak resistances to correct the current evoked by the sinusoidal protocol in both vehicle and dofetilide conditions. We then applied this leak resistance to the remaining protocols performed in the same condition on each cell.

The mean current during the –80 mV step was calculated from 200 ms of the –80 mV holding period before the –120 mV leak step (not including the capacitive spike at the point at which the step occurs). The baseline current at a holding potential of –80 mV was then adjusted back to 0nA with an additional constant additive current if required.

### 2.3 Dofetilide Subtraction

In preliminary work, we observed that our sinusoidal protocols could elicit endogenous voltage-dependent background currents within expression-system cells. We observed that the levels of endogenous currents the protocols elicited varied from cell to cell. These currents could adversely affect the predictive ability of the resulting mathematical models, as the fitting process attempted to create a model that described both the endogenous and I_Kr_ components of the recorded currents. To overcome this technical issue we made a number of alterations to our pilot experiments.

Firstly, we constrained the design of the sinusoidal protocol, as discussed above, so that only voltages within a physiological range of –120 mV to +60 mV were explored, as endogenous currents were much more prominent at voltages above +60 mV that we explored in pilot studies.

Secondly, we changed to use CHO cells in this study, rather than the HEK cells we used in pilot studies, as CHO cells generally had lower endogenous currents.

Thirdly, we recorded the full set of voltage protocols (Pr1–7) twice: once in Dimethyl sulfoxide (DMSO) vehicle conditions and once following the addition of 0.3 *μ*M dofetilide, as shown in Figure 2. Dofetilide was first dissolved in DMSO before being added to the bath solution to produce the required concentration. The required dose of dofetilide was obtained by serial dilution. We chose to use 0.3 *μ*M because the dofetilide hERG IC_50_ value is <10 nM which, assuming a Hill coefficient of one, should correspond to >97% conductance block of I_Kr_ at 0.3 *μ*M dofetilide. We avoided higher concentrations as dofetilide does have other known voltage-dependent ion channel targets whose IC_50_s are in the 10s–100s of *μ*M range (Mirams et al., 2011). Between the two recordings we allowed the dofetilide-induced current block to reach equilibrium (under Pr0). We then subtracted the currents that remained in the presence of dofetilide from those recorded in the presence of vehicle to remove any contribution of endogenous currents (and to produce what we refer to as ‘dofetilide subtracted’ data). Prior to performing this subtraction, we first leak subtracted both the vehicle and dofetilide recordings individually, as described above. It may not always be necessary for dofetilide subtraction to be performed on CHO cells, as endogenous voltage-dependent currents can be very low, and leak subtraction may suffice (see Appendix B1.6). But we applied the dofetilide subtraction method nonetheless to generate a gold-standard dataset for this study.

### 2.4 Mathematical Model

Whilst our model is equivalent to a two gate Hodgkin-Huxley formulation, we use a Markov model description in practice (simply to generalise the computational code for other model structures; the relationship between equivalent Markov and Hodgkin-Huxley models is explained in Keener and Sneyd (2009), vol. 1, p150). The system of ordinary differential equations underlying the mathematical model structure shown in Figure 4B is then:

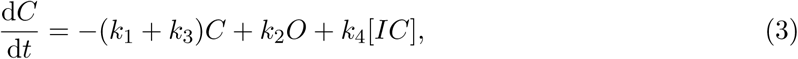

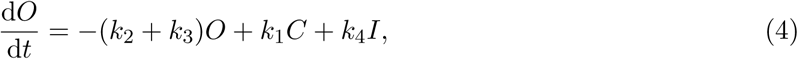

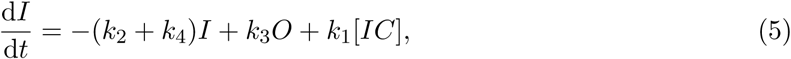

where the fourth state is constrained by probabilities of state occupancies summing to one

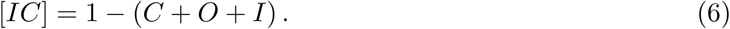

**Figure 4:**
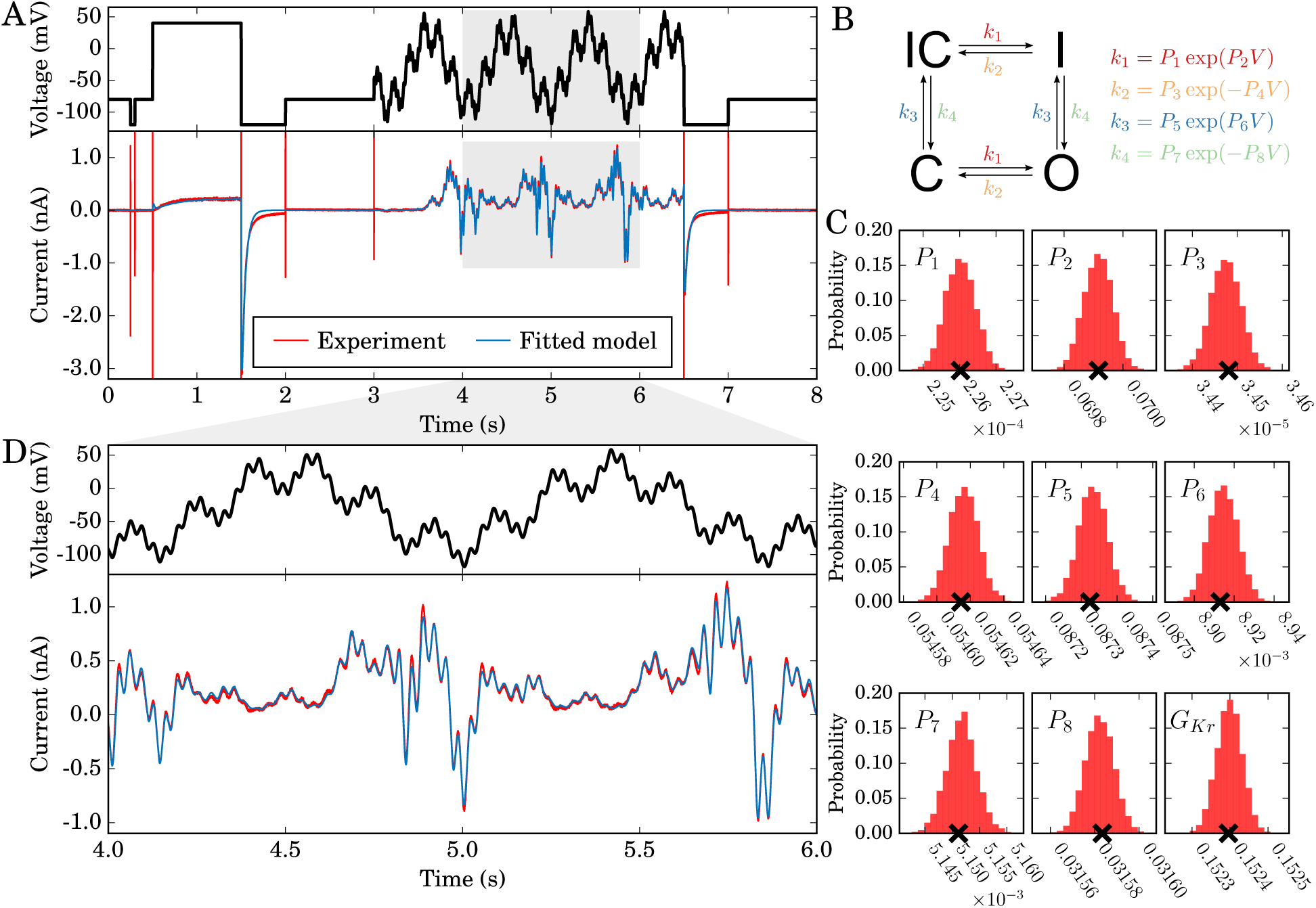
Model calibration. **A:** Top: the entire 8 second training protocol, bottom: an experimental recording with the fitted model simulation overlaid (portion of the sinusoid enlarged in panel D). This simulation uses the maximum posterior density parameter set, denoted with crosses in panel C. **B:** The model structure in Markov state diagram format, note that the symmetric transition rates mean this is equivalent to a Hodgkin and Huxley-style model with two independent gates. Parameter values *P*_1_ to *P*_8_ define voltage (*V*)-dependent transitions (*k*) between conforma-tional states. **C:** posterior distribution of single-cell derived model parameters. Probability density distributions are shown for each parameter after fitting to the experimental data shown in panel A. The parameter numbering corresponds to that shown in panel B. Crosses indicate the parameter set with the maximum posterior density. The standard deviation of each of these distributions is less than 0.2% of the maximum posterior density value. **D:** an enlargement of the highlighted region of panel A.

The eight parameters *P*_1_ to *P*_8_ determine the rates *k*_1_ to *k*_4_ according to the exponential voltage-dependence relationships shown in Figure 4B. The current, *I*_*Kr*_, is modelled with a standard Ohmic expression:

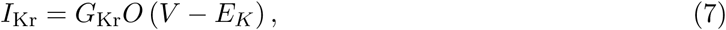

where *G*_Kr_ is the maximal conductance, *E*_*K*_ is the Nernst potential for potassium ions, and *O* is the open probability, given by the solution to the system of equations above. *E*_*K*_ is not inferred, but is calculated directly from the ratio of ion concentrations on each side of the cell membrane using the Nernst equation:

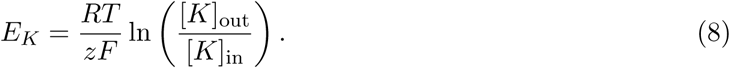

where *R* is the ideal gas constant, *T* is the temperature, *F* is the Faraday constant, z is the valency of the ions (in this case 1), and [*K*] represents the concentration of potassium ions. Note that this expression has a temperature dependence, and the temperature of the bath was recorded for each cell and used in relevant simulations.

All simulations were performed in MatLab. Mex functions were used to define the equations and simulate by using CVODE (Hindmarsh et al., 2005) to solve the systems of differential equations, with both absolute and relative tolerances set to 10^−8^. Code is available to download as described at the end of the manuscript.

### 2.5 Parameter Inference

We used a global minimisation algorithm (Hansen et al., 2003) followed by a custom-written Bayesian inference method. Parameters were estimated using a Monte Carlo based inference scheme, in this case using an approach similar to that described in Johnstone et al. (2016). In Appendix B2 we give details of how: (1) a likelihood is assigned to any candidate parameter set; (2) maximising the likelihood using a global optimisation scheme gives a ‘best fit’ parameter set; (3) uniform prior distributions are assigned to the kinetic parameters; and (4) we start a Markov chain Bayesian inference scheme from the estimated global optimum to generate a posterior probability distribution. The benefits of this scheme are that we explore the ‘parameter space’ widely and build up a probability distribution (probability of parameters generating the experimental results we observed) across the whole parameter space, thereby characterising any uncertainty in the ‘best fit’ parameter set. This posterior distribution allows us to check that we are constraining each parameter’s value with the information in the experiment, and are not experiencing problems with identifiability of parameters (Siekmann et al., 2012).

### 2.6 Note on Normalisation

Where existing literature model simulations were plotted alongside experimental traces, or one experimental trace was compared with another, we first had to normalise to account for differences in conductance values. This was achieved by selecting a scaling factor for the conductance value for each model simulation (or experimental trace) that minimised the square difference between each trace and a reference experimental trace.

For literature models the reference trace was the experimental current from the action potential clamp Pr6. Note this provides a best-case fit to Pr6 for all of the literature models, removing the possibility that some models open ‘half as much’ because they have ‘twice the conductance’. For the new model, no scaling was applied and conductance was directly fitted to the experimental current from the sinusoidal protocol (along with other parameters).

## 3 Results

### 3.1 Model Calibration

We calibrate a mathematical model using only data recorded under the sinusoidal protocol (Pr7). The Hodgkin and Huxley-style structure of the model we use, and its corresponding model parameters, can be seen in Figure 4B. We independently fitted this model to each of the experimental current traces shown in Figure 3. For each cell, we obtain a probability distribution of estimates for each parameter that captures any observational uncertainty in the parameter values (Pathmanathan and Gray, 2013; Mirams et al., 2016).

The result of the fitting procedure for one cell is shown in Figure 4. The parameter set with maximum posterior density is shown in Figure 4A, demonstrating an excellent fit between experimental and simulated data. The resulting posterior probability density for the parameters obtained from this Bayesian inference approach is projected across each parameter in Figure 4C. We also tested that our approach is theoretically appropriate for inferring all parameters by using synthetic data studies, as described in Appendix C. The plausible parameter space is very narrow: if multiple parameter set samples are taken from the distribution shown in Figure 4C, the resulting simulated current traces are indistinguishable to the eye. To quantify this, taking 1000 samples we find that the 95% credible intervals for the simulated currents were always within at most either 3.47% or, in absolute terms, 0.0043 nA of the simulated current given by the maximum posterior density parameter set.

The results we present in Figure 4 are from a single cell with a good quality recording and a high signal:noise ratio (this choice of cell, and other cells’ predictions, are discussed later). We fit models on a cell-specific basis, and then also use averaged experimental data to create a single ‘averaged’ model as described in Appendix F. We will compare these approaches below. We provide all parameter values with the maximum posterior density for all models in Appendix Table F11.

### 3.2 Validation predictions

Having trained our model to eight seconds of experimental data from the sinusoidal protocol Pr7, we now test its ability to predict more than 5 minutes of independent experimental behaviour. We predict the current in response to traditional voltage-step protocols Pr1–5 (adapted from those previously used in the literature (Bett et al., 2011)), and also to a novel physiologically-inspired voltage clamp protocol comprised of multiple action potentials (Pr6). All recordings shown in Figures 4–6 are from the same cell, using the experimental procedure shown in Figure 2.

To make the predictions for Protocols Pr1–6 we performed simulations using the parameter set with the maximum posterior density in the fit to the sinusoidal protocol (Pr7). As with the calibration protocol, all the predictions we will discuss below are indistinguishable by eye from the result of taking multiple samples from the distributions in Figure 4C and plotting a prediction for each of these parameter sets.

In Figure 5, we show traditional voltage step protocols, experimental recordings and the simulated predictions from the model. We also present some of the most commonly-plotted summary curves for experimental data under these protocols, together with predicted summary curves from our model. We compare these results with the summary curve predictions from a sample of widely-used literature models. We chose models for hERG1a expression systems at room temperature (Wang et al., 1997; Di Veroli et al., 2013) and physiological temperature (Mazhari et al., 2001); and also models with the same Hodgkin-Huxley structure as ours (Zeng et al., 1995; Ten Tusscher et al., 2004) albeit for physiological temperatures, as these are most directly comparable (methods used to derive summary plots are given in Appendix B1.7 with some additional summary curves for Pr1, 2 & 4 in E). We can predict a wide range of current behaviour in response to the standard voltage-step protocols, without having used any of this information to fit the model.

**Figure 5:**
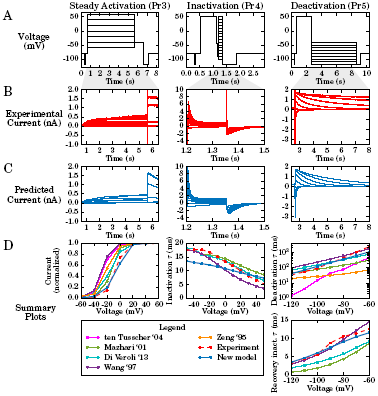
Validation predictions — currents in response to traditional voltage step protocols. Each column of graphs corresponds to a validation step protocol: those commonly used to study steady state activation, inactivation and deactivation (Pr3, Pr4, Pr5 in Fig 3) respectively. **A:** the voltage protocols. **B:** experimental current traces. **C:** model response — all are predictions using the maximum posterior density parameter set indicated in Fig 4C calibrated to just the sinusoidal protocol. **D:** summary curves, either current-voltage (I-V) or time constant–voltage (*τ*-V) relationships. These plots summarise the results in the relevant column. The model prediction is shown in blue bold throughout, and the experimental recording with a dashed red line. Note that the deactivation time constant we plot here is a weighted tau, described in Methods B1.7. Note that some literature model predictions are missing from the summary plots as we were either unable to fit exponential curves to ‘flat’ simulation output reliably; or the exponential decay occurred in the opposite direction to experimental traces, and we considered the comparison unwarranted.

**Figure 6:**
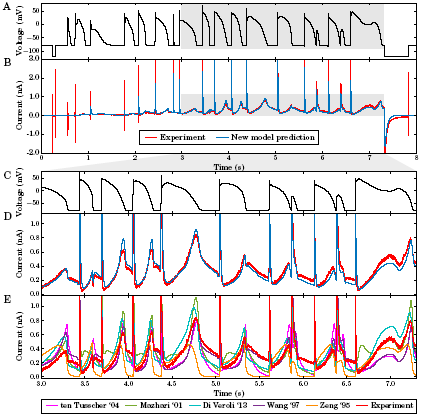
Validation prediction — the current in response to the action potential protocol. **A:** the voltage clamp protocol. **B:** a comparison of the experimental recording (red) and new model prediction (blue). **C & D:** enlargements of the highlighted regions of panels A & B. **E:** the same view of the experimental data in panel D, but here compared with predictions from literature I_Kr_ models. Conductance, *G*_Kr_, is scaled for each of the literature models to give the least square difference between their prediction and these experimental data, i.e. we display a best-case scaling for each of these models. A quantification of the error in our model prediction versus these literature models is given in Appendix Table D6: the performance shown in panels D and E holds for the whole trace, so the mean error in predicted current across the whole protocol is between 69% and 264% larger for the literature models’ predictions than for our sine-wave fitted model.

There are a number of points to draw attention to in Figure 5. Firstly, most of the current-voltage relationships and time constant-voltage relationships we predict in response to the traditional voltage-step protocols are closer to the experimental data than similar model-experiment comparisons in the literature (even when existing literature models, with more parameters, were fitted to such data). Secondly, there are some weaknesses to the new model — particularly in predictions of the Pr4 summary plot of time constant (*τ*) of inactivation against voltage, where we predict a time constant that is approximately 4 ms too fast at –40 mV. Yet, it is worth noting that this may be the best fit that is possible with a Hodgkin-Huxley style model: the Ten Tusscher and Zeng models predict timecourses that are so different it is difficult to fit comparable time constants. The current timecourse for Pr4 is actually predicted more accurately than any of the other models shown here (see Appendix Table D6) despite the *τ*-V relationship being less accurate; in agreement with this, other summary IV curves of Pr4 are predicted more accurately by the new model (see Appendix Figures E9 & E10).

Figure 6 shows the model prediction of the currents invoked in response to the physiologically-inspired action potential protocol Pr6, compared with the experimental recording (as shown in Figure 2 we used the first repeat of Pr6 for validation purposes, and the second as a quality control measure). Replicating behaviour under action potentials is perhaps the most important requirement for a hERG channel model for use in physiological or pharmacological studies. The model is able to predict the response to all of the complex action potential protocol extremely well, and much better than existing models (even though we have scaled all the literature models’ maximal conductances (*G*_Kr_) to fit this trace as well as possible in Figure 6).

We provide a quantitative comparison of predicted current traces for our model and each of the literature models for Pr3–7 in Appendix Table D6. In each case, the worst-performing literature model is a Hodgkin-Huxley style model. Yet our simple model, with the same structure, is able to provide significantly better predictions than even the Markov-type models, which are usually considered to be better representations of hERG kinetics (Bett et al., 2011). Our methodology has resulted in a simple and highly predictive mathematical model, able to describe a wide range of physiologically-relevant behaviour.

#### 3.2.1 Cell-specific validation

In Figure 7A we present the maximum posterior density parameter values when repeating the above approach using data from nine different cells. The clustered parameter values demonstrate that parameters derived from different cells take similar values, giving us confidence that the procedure is reproducible and biophysically meaningful. There is more cell-to-cell variability in some parameters than others, which may be related to variability in the underlying physiological processes that they represent; supporting the value, and perhaps necessity, of a cell-specific approach. We also acknowledge that some parameters may be more or less sensitive to variability in experimental conditions such as temperature, residual background/endogenous currents, and imperfect dofetilide and/or leak subtraction.

**Figure 7:**
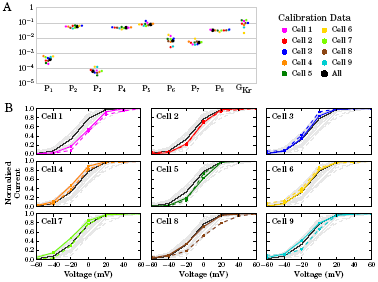
Cell-specific model parameters, and comparison of their predictions with cell-specific experimental results. (**A:**) Plot of parameters (maximum posterior density values) for nine cells obtained from training the model to the sinusoidal voltage protocol recorded on nine different cells, together with parameters calibrated to average data (N.B. not the average of the cell-specific parameters). The full set of parameter values are shown in Appendix Table Fff and the distributions for each parameter shown in Fig Fff. (**B:**) Comparison of cell-specific model predictions to cell-specific experimental recordings for the steady-state peak current I-V curves from Pr3. Each plot represents a different cell, model predictions are depicted by a bold coloured line, and dashed lines show values derived from the experimental data. The black lines (same on each plot) represent the prediction from the model calibrated to averaged sinusoidal data (all of the cells’ data). Each subplot contains all of the other cells’ recordings and predictions in light grey in the background to aid comparison and show the spread that we observed.

We order the cells in Figure 7 based on the lowest to highest difference in leak resistance between the vehicle and dofetilide recordings of Pr7. This ordering gives a measure of recording stability, and is intended to be a surrogate for data quality. The cell presented above, in Figures 4–6, corresponds to Cell #5 of 9 under this ranking, so we obtain very good predictions even with our ‘median’ quality data. We show cell-specific predictions of the current-voltage relationship for the peak steady-state activation current for each cell-specific model in Figure 7B. While we focused on Cell #5 in the results section, Cells #1–4 also produce excellent cell-specific predictions (similar comparisons for other summary plots are in Appendix Figures F12–F14).

We also investigated the benefit of our cell-specific approach by building a model using averaged experimental data from all nine cells instead. We describe this approach in Appendix F, and summarise the results in Appendix Table F12. Generally, for the cells with the highest data quality (Cells #1–5) the cell-specific models provide better predictions than the average model, as we see for Pr3 when comparing coloured cell-specific predictions and experiment with the black line for the average model in Figure 7B. The same trend holds for the action potential protocol Pr6, in 8/9 cells the cell-specific model provides less error than the average cell model — the largest improvement was 50% less error; for the remaining cell where the average cell model provided better predictions, this was by 3%.

## 4 Discussion

In this paper we have presented a novel method for capturing ion current properties, based on constructing mathematical models of ion channel kinetics. We used a sinusoidal voltage protocol to construct a simple model of hERG channel kinetics using just 8 seconds of recording, as opposed to a traditional approach that requires several minutes of voltage-step data. All of our experimental data can be collected from a single cell; whereas traditional protocols require long experiments, and typically require different gating processes to be studied in different experiments in different cells. In future, our approach opens up the possibility of making multiple interventions (such as the addition of drug compounds) since we could re-measure the full ion channel kinetics multiple times in a single cell.

The conceptual shift is that channel kinetics should be summarised by mathematical model parameters, not a series of current-voltage (IV) and time constant-voltage curves. In essence, the model is the current characterisation, rather than something designed to fit IV and time constant curves, which only represent a certain subset of possible behaviours of the current. The success of the approach lies in moving away from traditional protocols that can be easily interpreted by eye, which typically require the current to return to an equilibrium rest state between voltage steps. Instead, our protocol also probes non-equilibrium ion channel behaviour by rapidly exploring time and voltage dependence, and is interpreted through the fitting of a model for the whole current at once.

Our model is able to replicate the experimental training data very well (Fig 4). This is often the point at which traditional approaches in the literature have stopped, and concluded that a mathematical model is a good representation of ion channel kinetics (also true more generally for mathematical models of biological processes). Instead, we performed an extremely thorough evaluation of the model by testing its ability to predict the behaviour in response to a series of voltage clamp protocols it has not ‘seen before’ (both those traditionally used to characterise hERG channel kinetics, and also a new complicated series of action potential waveforms), all recorded from the same cell as the training data. We are not aware of such a thorough, physiologically-relevant validation of an ion channel model having been performed before. Testing that we are able to predict the current response to a voltage pattern which may be observed in physiological or patho-physiological conditions is a particularly robust and useful way to validate a model, and critical if an I_Kr_ model is to be used to accurately predict cardiac electrical activity in both healthy and potentially arrhythmic situations.

The extremely good prediction from all our cell-specific models of the response to the complex action potential protocol is particularly remarkable (Fig 6). Cell-to-cell variability in ion channel kinetics was captured by fitting different underlying kinetic parameters. These parameter sets were shown to have modest variation, and this variation in kinetics was quantitatively predictive of variation observed in independent validation experiments (Fig 7).

Cell-specific predictions were particularly strong when using the highest quality data, highlighting the necessary data quality for constructing accurate and robust models of ion channel kinetics. The cell-specific models outperformed a model constructed using averaged data from multiple cells/experiments, in line with the ‘failure of averaging’ discussed in Golowasch et al. (2002) and the problems of fitting to averaged summary curves outlined in Pathmanathan et al. (2015).

Our inactivation protocol (Pr4) showed that it is possible for models to fit some (or all) summary curves well, without necessarily replicating the underlying current traces with less error. Often studies present just single summary curves in isolation. But we have seen how models can fit certain summary curves well, whilst fitting others badly. Models that have less accurate summary curves may even predict the underlying current traces more reliably; and, importantly, vice-versa. A focus on these summary curves to represent kinetics and fit mathematical model behaviour was necessary in the era of hand fitting parameters using graph paper, but should perhaps now be superseded by fitting/comparing directly to experimental current traces. By fitting directly, we also reduce the possible influence of subjective choices during time-constant fitting used in the generation of time-constant voltage relationships.

A limitation of our study is that our model was trained on experiments performed in expression line cells, creating a hERG1a model at room temperature; compared to native I_Kr_ current in cardiac cells which will have additional isoforms, subunits and regulation at physiological temperatures. As a result, we do not state that this ion current model would necessarily give better performance within a cardiac action potential model. To characterise native I_Kr_ kinetics we plan to apply the methodology presented here in myocytes, to make a model that is more applicable for use in cardiac safety testing and whole-organ simulations. The presence of many larger voltage-dependent currents than we observe in expression systems will make this challenging, but a dofetilide subtraction approach may still yield good results.

There are still some aspects of the experimental behaviour that are not replicated by our model. These aspects may be a consequence of using a simple Hodgkin-Huxley style model formulation, although it remains a commonly-used structure for currents within action potential models. In particular, there is only one time constant of deactivation, and low voltage-dependence in the inactivation time constant (Fig 5). A more complicated model with additional states and parameters may be needed to capture certain behaviours.

We assessed the capability of the protocol to fit a more complex five state Markov model for hERG (the model proposed by Wang et al., 1997), and show the results in Appendix H. Previously, Bett et al. (2011) explored the behaviours of a subset of existing hERG models and concluded that this model was best able to replicate activation kinetics. In Appendix H we show that exactly the same approach and algorithms again tightly constrained all 15 of the parameters in this larger model, using the same sinusoidal protocol data. The more complex model resulted in a better fit to the calibration data, and also made good predictions for the validation protocols — although not quite as good as the simpler model presented here in the main text. This finding highlights the importance and challenges of selecting the most appropriate level of complexity for a mathematical model.

So despite our simple model not replicating precisely the full range of behaviour, neither do the existing, more complex, models available in the literature. We have shown that our simple model can provide better predictions than the literature models for all the raw current timecourses, if not all summary curves, in the majority of cells. In fact, the simplicity of our model may be the key to its success — with only eight kinetic parameters we have confidence that they are all being fitted well, and we have shown that there is low uncertainty in their values.

The applicability of our approach for different ion channels will be heavily dependent on the precise form of the sinusoidal protocol that is used, and in parallel work we are developing different strategies for optimising the voltage protocol design for given currents. Although we have also shown that the existing protocol is at least theoretically appropriate for parameterising an I_Ks_ model in Appendix G. In future work, ideas from control engineering may be useful. Seemingly unconnected problems, such as generating signals to characterise the state of lithium ion batteries (Xiong et al., 2011), are in fact very similar mathematical challenges.

There will be limits in the complexity of model structure and number of parameters that any protocol can constrain. But in terms of limitations of this style of protocol, we consider that the more information-rich protocols are, the better; and these new protocols may enable us to accurately calibrate larger models than before. We strongly advocate synthetic data studies to assess the suitability of a given protocol for constraining parameters of a given model — seeing whether re-fitting to data generated by simulations of your model and protocol can recover the parameters used in the simulation. Such approaches are necessary but not sufficient: they still rely on the models being a good representation of the system under study, and incorporating statistical ideas to handle model discrepancy (the difference between models and reality) is an important line of enquiry (Strong et al., 2012). In other parallel work, we are extending the approach presented here for selecting between different possible model structures for hERG channel kinetics (see Appendix Fig A1 for the range of possibilities; and Kargol (2013) for an outline of how this may be approached by optimising the protocols themselves to assist with this task).

Considering probabilistic uncertainty in model parameters and predictions is evermore important as models begin to be used for safety-critical predictions (Pathmanathan and Gray, 2013; Mirams et al., 2016). These predictions include guiding therapies (Arevalo et al., 2016) and pharmaceutical safety assessment with the Comprehensive in-vitro Proarrhythmia Assay initiative being pursued by the FDA in collaboration with industry, academia and other regulators (Sager et al., 2014; Fermini et al., 2016). Here we have shown there is very low uncertainty in hERG kinetics parameters in a single cell, and also characterised the variability in these estimates between different cells.

In summary, we have demonstrated significant advantages in our cell-specific mathematical modelling approach, observing excellent model predictions of currents in response to protocols the model was not trained to replicate. The simple ion channel state arrangement we have assumed must capture the most important features underlying hERG state transitions, despite being much simpler than many previous models in the literature. The information-rich approach allows, for perhaps the first time, an exploration of both within-cell and between-cell variability in ion channel kinetics. The significant time saving of our short protocol also leads to datasets that are more consistent and therefore of higher quality, since little changes in experimental conditions during the 8 second recording interval. Its brevity opens up the possibility of taking more recordings in different experimental conditions within a single cell (e.g. drug concentrations (Pearlstein et al., 2016; Lee et al., 2016) or temperatures (Vandenberg et al., 2006)). These datasets will result in more accurate descriptions of ionic currents in these different conditions in the heart and other organ systems.

## Acknowledgements

Our thanks to Prof. Gail Robertson of University of Wisconsin–Madison for assistance in acquiring the cell line used in pilot stages of this study. We would also like to thank the following people for technical assistance, access to facilities, support and encouragement: Jim Louttit, Nick McMahon, Carol Wilson, Sam Turner, Kate Harris and Sara Graham of GSK Safety Assessment; Jules Hancox of University of Bristol; Monique Windley and Mark Hunter of Victor Chang Cardiac Research Institute; Rianne Rijken and Birgit Goversen of UMC Utrecht. Thanks to Ross Johnstone (University of Oxford) for removing singularities from the Zeng model, and to Frank Ball (University of Nottingham) for comments on a paper draft.

## Author Contributions

KAB, RB, YC, DJG, TdeB, GRM designed the study and modelling approach; KAB, RB & GRM designed and implemented the statistical methods; KAB, GRM, JIV, APH and TdeB designed and refined the experimental methods; KAB performed all the experiments, simulations and statistical analysis; KAB, TdeB, GRM wrote the manuscript; all authors approved the final version of the manuscript.

## Materials

All computational codes, and the experimental current recordings that were used for calibration and validation (leak and dofetilide subtracted), are openly available in a Supplementary Data repository at https://github.com/mirams/sine-wave. A permanently archived version is available on Figshare at https://doi.org/10.6084/m9.figshare.4704550.v5 alongside the full raw data (in both plain text and pClamp formats) at https://doi.org/10.6084/m9.figshare.4702546.v1.

For additional details on the methods please see the online Appendix.

## Footnotes

1 This article was first published as a preprint on bioR*χ*iv: Beattie *et al.* (2017): https://doi.org/10.1101/100677

## Non-standard Abbreviations and Acronyms

CHO: Chinese Hamster Ovary [cells].
G_Kr_: maximal conductance of I_Kr_.
HEK: Human Embryonic Kidney [cells].
hERG: human Ether-a-go-go Related Gene.
I_Kr_: rapid delayed rectifying potassium current, carried by the K_v_11.1 ion channel whose primary subunit is encoded by hERG.

